# Host and Water Microbiota are Differentially Linked to Potential Human Pathogen Accumulation in Oysters

**DOI:** 10.1101/2022.10.11.511793

**Authors:** Rachel E. Diner, Amy Zimmer-Faust, Emily Cooksey, Sarah Allard, Sho M. Kodera, Emily Kunselman, Yash Garodia, Andrew E. Allen, John Griffith, Jack A. Gilbert

## Abstract

Oysters play an important role in coastal ecology and are a globally popular seafood source. However, their filter feeding lifestyle enables coastal pathogens, toxins, and pollutants to accumulate in their tissues, potentially endangering human health. For example, bacterial pathogens from both marine and terrestrial sources concentrate in oysters and can cause human illness when oysters are consumed raw. While pathogen concentrations in coastal waters are often linked to environmental conditions and runoff events, these do not always correlate with pathogen concentrations in oysters. Additional factors related to oyster hosts and the microbial ecology of pathogenic bacteria likely play a role in accumulation but are poorly understood. In this study, we investigated whether microbial communities in water and oysters were linked to accumulation of fecal indicators, *Vibrio parahaemolyticus*, and *Vibrio vulnificus*. Site-specific environmental conditions significantly influenced the composition and diversity of water microbial communities, which were linked to the highest concentrations of both *Vibrio* spp. and fecal indicator bacteria. Oyster microbial communities, however, were less impacted by environmental variability and exhibited less variability in microbial community diversity and accumulation of target bacteria. Instead, changes in specific microbial taxa in oyster and water samples, particularly in oyster digestive glands, were linked to elevated potential pathogens in oysters, especially *V. parahaemolyticus*. This included an increase in cyanobacteria in both water and oyster digestive gland microbial communities, which could represent an environmental vector for *Vibrio* spp. transport and decreased relative abundance of *Mycoplasma* and other key members of the oyster digestive gland microbiota. These findings suggest that host and microbial factors, in addition to environmental variables, may influence pathogen accumulation in oysters.

## Introduction

Bivalves are ecologically important animals that provide valuable coastal services and are a globally popular, potentially sustainable food source. In addition to recreational and subsistence harvesting, bivalve aquaculture operations generate numerous local jobs and approximately US$23 billion in revenue annually (1). Oysters and mussels build coastal reefs which provide habitat for coastal organisms and can prevent erosion (2, 3). Additionally, adult bivalves filter large quantities of water to concentrate their phytoplankton prey, which can reduce nutrient loads and improve coastal water quality (3).

The ability of bivalves to thrive in highly fluctuating coastal environments can be impacted by human activities and environmental stress, impairing the provision of these ecosystem services and their commercial potential. Additionally, while concentrating their prey, many bivalves concentrate human pathogens, marine toxins, and coastal pollutants, which can endanger human health when consumed. These risks to seafood safety and security are likely to increase in the future due to global changes such as climate change, increasing human populations, pollution discharge, and rapid coastal development.

Bacterial pathogens are a major food safety concern, especially for oysters which are commonly consumed raw. Infections can be caused by fecal-borne bacteria from terrestrial sources as well as marine species, which thrive in the oyster’s natural environment. For example, *Vibrio* species cause an estimated 80,000 human illnesses in the US annually and 100 deaths (4). Among these, *Vibrio parahaemolyticus* is the most common cause of infection, while *V. vulnificus* causes the greatest mortality, with 1 in 5 cases resulting in death. As these species thrive in warm conditions, they are highly concerning amidst rising global seawater temperatures (5, 6). Additionally, since *Vibrio* spp. naturally occur in marine and brackish waters, they may resist depuration processes commonly employed to remove pathogens from oysters prior to entering the food supply. Understanding the factors that cause bacterial pathogens to accumulate in bivalves is critical to preserving human and ecosystem health now and in the future.

The environmental drivers of human pathogen concentrations in coastal waters are relatively well-characterized, however, the connection between water and oyster pathogen concentrations is less clear. For fecal-borne bacteria, high concentrations in coastal waters are often linked to freshwater influx and sewage spills, frequently coinciding with high concentrations of nutrients and other pollutants. Marine *Vibrio* species are associated with warm water temperatures and exhibit species-specific salinity distributions (reviewed in (7, 8)). They also attach readily to marine particles which can enhance oyster accumulation. While the presence of pathogenic bacterial taxa in the water may be a prerequisite for oyster accumulation, studies investigating water to oyster pathogen transfer are either rare or yield inconsistent results, suggesting that factors beyond the environment likely play a role. For example, individual oysters from the same environment can have highly variable pathogenic bacterial concentrations, known as the “hot oyster” phenomenon (9, 10), and associations between water and oyster concentrations of pathogenic *Vibrio* species vary among studies and geographic regions. Ultimately, more research is needed to examine different variables that could explain the relationships between environmental factors, water to oyster human pathogen transfer, and pathogen accumulation and persistence in oysters.

Microbial communities associated with water and oysters are understudied variables that likely influence human pathogen accumulation. Microbial taxa in the water column may help pathogenic species proliferate in the environment and act as vectors for uptake and concentration by shellfish. For example, *Vibrio* species have been linked to phytoplankton concentrations in coastal waters, and human pathogenic species have shown associations with specific phytoplankton groups and individual taxa (11–15). Several of these taxa are known oyster prey species. Furthermore, microbial communities associated with oyster hosts (i.e., their microbiomes) could also influence pathogen accumulation if environmental factors cause changes in otherwise stable microbial communities. Marine animal microbiomes often contribute to host health; critical functions include food breakdown, defense against host-associated pathogens, and modulation of host immune responses (Reviewed in (16–18)). In stressful conditions, however, animal microbiota can shift from a health-associated state, to “dysbiosis”, whereby the microbiota and host no longer express beneficial synergy. This often leads to proportional increases in environmental microbes that are typically rare or absent in host microbiomes, or an increase in commensal host microbiota that can become pathogenic. In aquaculture organisms, host-specific pathogens are well studied in this context, but the accumulation of human pathogens in animal hosts as a function of dysbiosis has not been adequately investigated.

To investigate the influence of these understudied factors on potential human pathogen accumulation in oysters, we examined fecal indicator bacteria, *V. parahaemolyticus*, and *V. vulnificus* concentrations in water and oysters across an environmental gradient in a southern California coastal bay over 4 weeks. We then applied a metabarcoding approach (16S amplicon sequencing) to characterize links between these target bacteria, water and oyster prokaryotic microbiomes, and environmental variables. In addition to better understanding how concentrations of these bacteria in water are linked to oyster concentrations, our study aimed to characterize how environmental variation influences oyster microbiomes and test whether altered host-states may be linked to higher bacterial accumulation. Additionally, we sought to characterize specific microbial taxa in water and oyster microbiomes that correlate with human pathogen accumulation in oysters, which could indicate environmental reservoirs or vectors for pathogen transmission.

## Materials and Methods

### Experimental design and sample collection

Approximately 1,200 Pacific oysters were collected over a three-day period (July 31-August 2, 2019) from Newport Bay, CA, then transported to holding tanks located at the Kerckhoff Marine Laboratory in Corona Del Mar, CA within four hours of harvesting. At the Kerckhoff Marine Laboratory, oysters were pooled and arranged on perforated stacked trays in four 282.7 m^3^ flow-through seawater tanks for 14 days. Seawater was first filtered through a sand filter at 15 – 20 gallons per minute and then further disinfected with a Classic UV 80-Watt Series light (Aqua Ultraviolet, Temecula, California, USA) before entering the holding tanks. Following the two-week hold time, oysters were deployed across 12 sites in Newport Bay, CA for six weeks (Figure 1). For purposes of this study, only weeks 0-4 were analyzed, however, data was collected for all 6 weeks (19). Roughly 100 oysters were deployed in 23-mm plastic mesh oyster bags at each site. At the time of oyster deployment (Week 0), 10-12 oysters were collected and processed from each holding tank to characterize composite post-depuration microbial communities in whole oysters (N=10-12) and oyster tissue samples (gills and digestive glands [DG], N=10-12 each), and water was collected at each deployment site. Thereafter, both water and oyster samples were collected at weeks 1, 2, and 4. Grab water samples were collected from each site, while oyster samples were collected as composite samples of 10 to 12 individual oysters from each site at each timepoint and for each tissue type. Water samples were stored on ice, while oysters were stored in coolers with ice packs, and transported to the Southern California Coastal Water Research Project (SCCWRP) for further processing.

**Figure 1:**
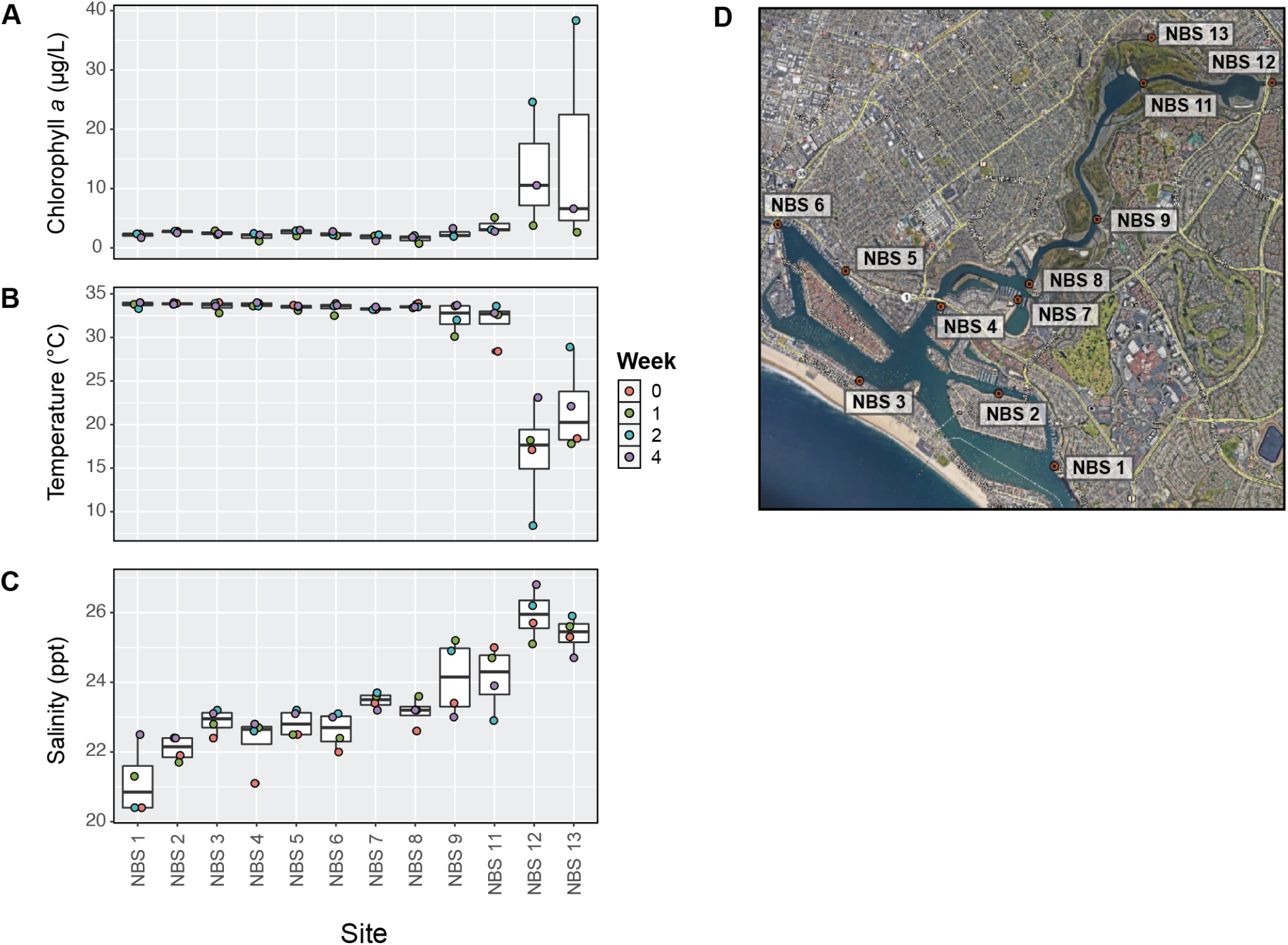
Experimental sites and environmental conditions. Environmental conditions at the time of sample collection, including (A) chlorophyll *a* concentration, (B) temperature, and (C) salinity. (D) Sites in Newport Bay, CA where oysters were deployed and water and oyster samples were collected for the study.

Oysters were collected at 12 different sites in Newport Bay, which corresponded to different attachment substrates depending on site topography, including dock, buoy, mudflat, and seawall (Additional File 2), though all oysters were ultimately translocated to identical cages following depuration. Oysters were not observed, and thus not initially collected, from sites NBS12 and NBS13 in the back bay. The environmental variations observed at these sites (Figure 1A-C) suggest that environmental or biotic factors may have made these sites inhospitable to endemic oyster growth, however low oyster populations could also be caused by illegal harvesting in these areas.

Water samples were collected in the field in cleaned and acid-rinsed (10% HCl) 2 L polycarbonate bottles. Samples were brought back to the laboratory after collection and were gently mixed before filtering. Approximately 100-200 mL of water was filtered onto 0.4 μm polycarbonate filters for nucleic acid extraction, which was used downstream to characterize water microbial communities. Additionally, water samples were collected to measure chlorophyll *a* concentration: 100 mL were collected on 25 mm glass fiber filters (Whatman) with gentle filtration and stored at −80 °C until processing. Lastly, more water aliquots were used to quantify fecal indicator microbial species, including *Escherichia coli*, total coliform bacteria, fecal coliform bacteria, and *Enterococcus* species. Filtered samples were stored in 1.5 mL tubes and frozen in Liquid nitrogen, followed by immediate storage at −80 °C until DNA extraction.

At SCCWRP, ~10 oysters were homogenized, and the composite was processed to quantify bacterial targets according to published methods (see below and Additional File 1). An additional ~10 oysters were shucked and dissected, and the gill and digestive gland tissues were separately pooled and homogenized. Homogenized digestive glands and gills were stored for downstream DNA extraction at −80 °C.

Environmental data was collected using a YSI30 Pro field instrument (Yellow Springs, OH), which was used primarily to assess temperature and salinity. Filters for chlorophyll *a* quantification were stored in aluminum foil in a plastic bag at −80 °C until further analysis, at which point they were extracted in 100% acetone in the dark for 24 hours. Total chlorophyll *a* was measured in all water samples using a non-acidification method and a Trilogy Turner Design Fluorometer, as previously described (20).

### *Quantification and detection of human pathogen and indicator species and OSHV-1* virus

The bacterial targets quantified in this study and the associated analysis methods are listed in Additional File 1. For bacteria enumeration in oyster tissues, we analyzed the fecal indicator bacteria (FIB) *Escherichia Coli* and fecal coliform, following approved methods developed by the FDA and Interstate Shellfish Sanitation Conference (Additional File 1). Briefly, fecal coliform and *E. coli* concentrations were determined by conventional five-tube multiple dilution most-probable number (MPN) procedure. Lauryl tryptose broth (Difco) was utilized for the presumptive growth media, with confirmation performed by inoculating liquid EC-MUG media (Difco) at 44.5 °C for 24 ± 2 hours. Grab water samples were processed for cultivable *Enterococcus, E. coli*, and fecal coliform according to standard methods: EPA Method 1600, EPA Method 1603, and SM 9222-D (21–23).

For *Vibrio* spp. targets, *Vibrio parahaemolyticus* and *V. vulnificus* were quantified using a culture based MPN method. Briefly, CHROMagar *Vibrio* (CHROMagar, Paris, France) media plates were prepared according to manufacturer’s instructions and used to enumerate potentially pathogenic *Vibrio* species. *V. parahaemolyticus* (Vp) and *V. vulnificus* (Vv) concentrations were determined by counting visible pink and blue colonies on the CHROMagar *Vibrio* media, respectively, and adjusting for dilution (24). Data was reported as CFU/100 mL and CFU/100 g for water and oysters and the limit of detection was 1 CFU/g or 1 CFU/mL. Up to ten presumptive Vp and Vv colonies per plate (when present) were stored for further species-level confirmation as described below. The number reported was then multiplied by percentage of molecularly confirmed (by PCR) isolates, resulting in confirmed bacterial abundance for each sample, as described previously (25).

To test for OsHV-1 presence and abundance, the ORF100 primer set described in Burge et al 2020 was used for qPCR (26). Briefly, 10 μL of PerfeCTa SYBR Green FastMix (Quantabio), 1 μL of each primer (ORF100 F, ORF100 R), 6 μL of water and 2 μL of DNA template were mixed per reaction. All samples were run in duplicate on the 96-well Agilent ARIAMx RT-PCR thermal cycler. A standard curve was generated with a synthetic plasmid of the ORF100 DNA sequence by serial dilution from 30 million copies down to 3 copies.

### Statistical Analysis of Environmental and Target Bacteria Variables

To determine whether the variable Site had a significant effect on concentrations of target bacteria and environmental conditions, non-parametric Kruskal-Wallis tests were conducted in R using the stats package. Pairwise comparisons were not conducted due to low sample number. To determine whether significant correlations existed between water and oyster samples for target bacteria, we conducted linear regression analyses. When data was not normally distributed as determined by the Shapiro-Wilk normality test, which occurred with Vp and Vv, a Tobit Regression analysis was conducted to determine significance of the observed correlation.

### DNA extraction, amplicon library preparation, and sequencing

DNA was extracted from water samples using the Qiagen PowerSoil kit, and from oyster tissues using the Qiagen DNEasy blood and tissue kit (Qiagen, Hilden, Germany). Digestive gland and gill composites, which were homogenized at the time of collection, were digested with Proteinase K prior to DNA extraction. The V4-V5 region of the 16S rRNA gene was amplified using 515F-926R primers (27). Libraries were assessed for quality using an Agilent 2200 TapeStation (Agilent, Santa Clara, CA). Since these primers also produce amplicons from 18S rRNA, BluePippin size selection was used to enrich for ~550bp amplicons prior to sequencing (Sage Science, Beverly, MA, USA). Additionally, 3 blanks (lysis buffer only, taken through extraction protocol, 2 extracted with tissue samples and 1 run with water samples) and 2 samples of DNA from mock bacterial communities (ZymoBIOMICS Microbial Community DNA Standard, Zymo Research, Irvine CA) were prepared and sequenced, with all samples included in the same sequencing run. After library construction, samples were sequenced using the Illumina MiSeq 2×300 (kit v.3) with custom adapters and dual barcode indices at the University of California Davis Genome Center (https://genomecenter.ucdavis.edu/). Raw data sequences and metadata are publicly available on the Qiita platform (28)(Study ID 14776) and the metadata file is published on figshare (https://doi.org/10.6084/m9.figshare.21272916).

### Sequence Quality filtering and Bioinformatic Analysis

The QIIME2 pipeline was used for quality control, filtering, and bioinformatic analysis (29). R packages, including Phyloseq (30), were used for additional analyses and data visualization. Demultiplexed sequences were imported as QIIME2 artifacts, and paired reads were trimmed to remove primers, merged, and assigned ASVs using dada2 (default parameters). Taxonomy was assigned to ASVs using the SILVA database (Version 138)(31). Prior to downstream analyses, mitochondrial, chloroplast, and eukaryotic sequences were removed as well as ASVs with no identified domain.

Diversity analyses were conducted in phyloseq without rarefaction, though rarefaction at a sampling depth of 2298 (determined by alpha rarefaction curves) was also compared to confirm that results were consistent between rarefied and non-rarefied datasets. Pairwise comparisons of alpha diversity were conducted in R using a Wilcoxon rank sum test, using the Holm method for P-value adjustment. For beta diversity analyses, Robust Atchison Principal Component Analyses (RPCA) were conducted and then visualized as biplots using the QIIME2 DEICODE plugin (32), with a minimum feature count of 10 and a minimum sample count of 500. PERMANOVA tests were used to analyze pairwise differences between groups (e.g., sample type, site) using the qiime diversity beta-group-significance command.

Log fold changes in the differential abundance of key taxonomic features in digestive gland samples were conducted in R by calculating the log ratio of targeted taxa of interest to core digestive gland microbial taxa in individual samples. Microbial taxa of interest were identified using three approaches (identification via DEICODE biplots, identification via taxonomy plots, and identification of ASVs belonging to Cyanobacteria) and analyzed separately. Core microbes were identified using the qiime feature-table core-features command, which identified 10 taxa present in >85% of samples. One of these taxa, one *Serratia* ASV, was removed from core microbiome consideration as it was also found in blank samples and is not considered a marine or oyster-associated bacterial taxa. For each sample, the abundances of taxa of interest were compared against abundances of core microbial taxa to provide differential abundance values. We used the following formula:

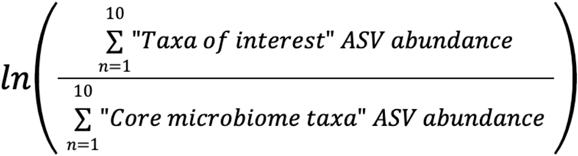

Correlations between metadata variables, including alpha diversity metrics, environmental variables, and pathogen concentrations were calculated based on Spearman’s rank correlations and visualized using the corrplot package in R (33). Additional correlations between these factors and the relative abundance of taxa of interest were conducted using data from weeks 0, 1, 2, 4 for water samples, and weeks 1, 2, and 4 from oyster samples since there was no week 0 for oysters at the designated sites based on the experimental design. Samples collected at the Kerchoff facility following depuration were not included in the correlation analyses.

## Results

### Environmental and oyster collection conditions at experimental sites

Experimental sites in Newport Bay, CA differed in environmental conditions including chlorophyll *a*, temperature, and salinity (Figure 1A-D, Additional File 2). Chlorophyll *a* concentration ranged from 0.72 μg/L to 38.34 μg/L (Figure 1A) and was significantly affected by site (Kruskal-Wallis rank sum test: chi-squared = 23.164, p-value = 0.017). In water samples, this metric is a common proxy for biomass of phytoplankton, which produce this photosynthetic pigment. Sites close to the back of the bay (i.e., NBS12 and NBS13) exhibited the highest observed concentrations, which occurred during weeks 2 and 3 (Figure 1A, Additional File 2). Temperature and salinity were also significantly affected by site (Temperature: chi-squared = 29.261, p-value = 0.002, Salinity: chi-squared = 23.598, p-value = 0.015). Salinity ranged from 8.4-34 ppt (Figure 1B) and temperature ranged from 20.4-26.8°C (67.7-80.2°F) across all sites (Figure 1C). Sites NBS12 and NBS13 exhibited warmer temperatures and lower salinities than other sites (Figure 1B, 1C).

### Target bacteria species concentrations in seawater and oysters

Fecal indicator bacteria (FIB) concentrations, which we quantified as a proxy for human fecal contamination, varied by site, and different abundance patterns were observed between water and homogenized whole oyster samples (“oyster samples”)(Figure 2, Additional File 3).

**Figure 2:**
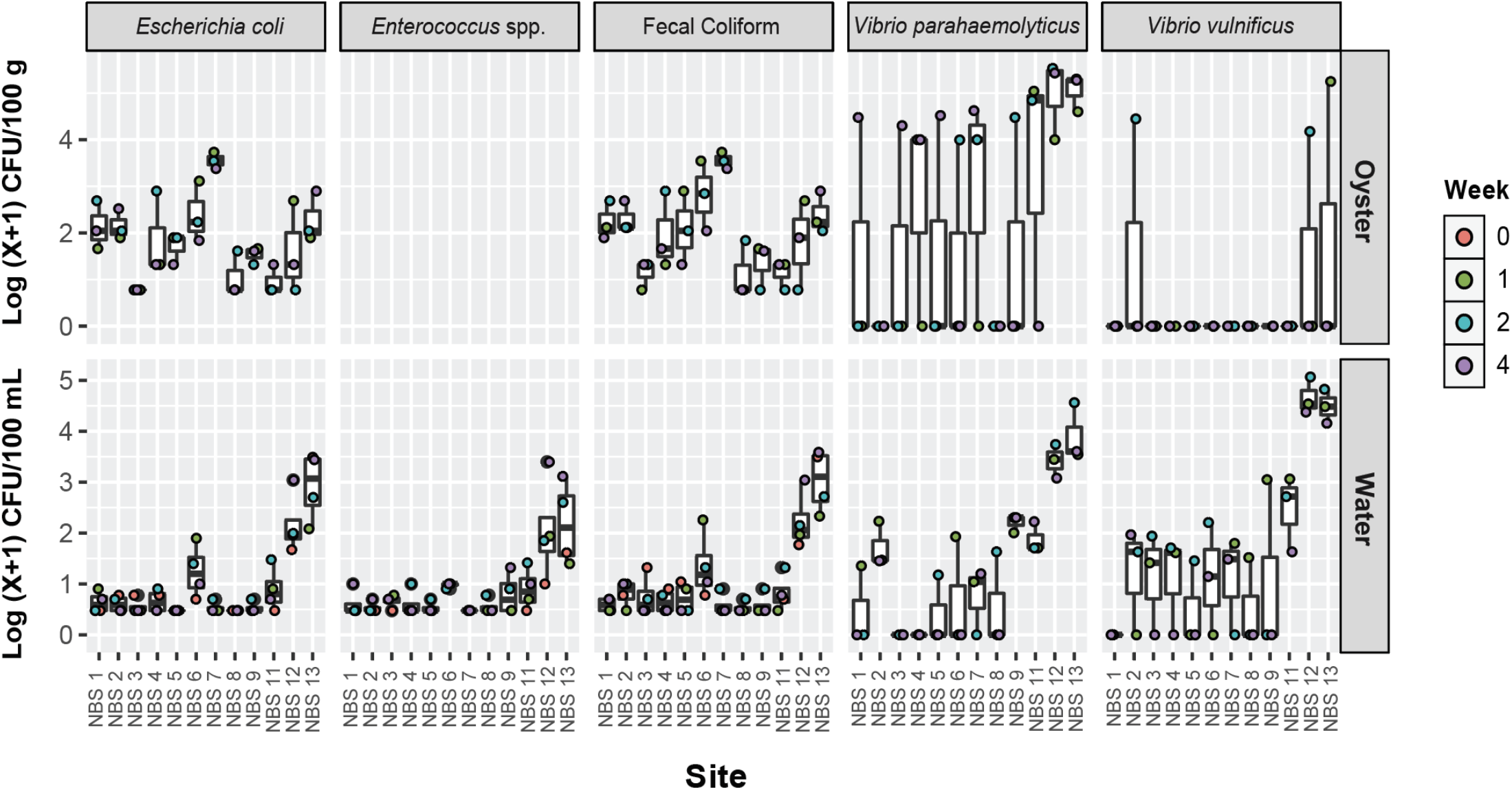
Fecal indicator bacteria (FIB) and pathogenic *Vibrio* spp. concentrations in oysters and water. Concentrations are shown as Log (x+1) CFU/100g of oyster tissue or 100 mL of seawater. FIB concentrations in oysters were calculated using the MPN method.

Concentrations of fecal coliform bacteria (FC), *Escherichia coli* (EC), and *Enterococcus* spp. (ENT) in water samples were all elevated at sites 12 and 13 compared to samples collected at other sites during the same week (Figure 2) and were overall significantly affected by site (FC: chi-squared = 23.476, p-value = 0.015, EC: chi-squared = 28.76, p-value = 0.002). FC and EC concentrations in water also appeared to be elevated at site 6, though positive samples were observed at all sites during the experiment. In oyster samples, FC and EC concentrations were elevated at site 7 compared to the other sites (Figure 2, Additional File 3) and were also significantly affected by site (FC: chi-squared = 23.921, p-value = 0.013, EC: chi-squared = 25.418, p-value = 0.008). Oyster and water concentrations were not correlated for *E. coli* and fecal coliforms (Linear Regression, E. coli: F-statistic: 0.6927, adjusted p-value: 0.411, fecal coliform: F-statistic: 1, adjusted p-value: 0.1916) (Additional File 3). Water column concentrations of FC and EC exceeded the California SHEL standard of 43 MPN/100 mL at 3 sites: NBS6, NBS12, and NBS13.

*Vibrio* species of concern for human health also varied in concentration by site but unlike with FIB, oyster and water samples were positively correlated, particularly for *Vibrio parahaemolyticus* (Vp) (Figure 2, Additional File 3). In water, *Vibrio* spp. targets were highest at sites 12 and 13 compared to different sites measured during the same week (Figure 2), which generally corresponded to warmer temperatures and lower salinities (Figure 1B, 1C). Maximum Vp concentrations exceeded 30,000 CFU/100 mL (Site NBS13, Week 2, Additional File 2) while highest *V. vulnificus* (Vv) concentrations were >100,000 CFU/100 mL (Site NBS12, Week 2). These species were also observed at most other sites during the experiment, albeit in lower concentrations. Site had a significant effect on water concentrations of both Vp and Vv (Vp: chi-squared = 29.351, p-value = 0.002, Vv: chi-squared = 22.371, p-value = 0.022). However, Site did not significantly affect Vp or Vv concentrations in oysters (Vp: chi-squared = 18.715, p-value = 0.066, Vv: chi-squared = 9.5502, p-value = 0.571). In oyster samples, Vp concentrations were also highest at sites 12 and 13 compared to other sites, except for during week 1 when concentrations were also elevated at site 11. Vv was only detected in 3 oyster samples, and the highest concentration (177,500 CFU/100 g oyster tissue) was observed during week 1 at site NBS13. Correlations between water and oyster concentrations were positive and significant for *V. parahaemolyticus* (Tobit Regression, Wald-statistic: 5.862, p-value: 0.015) and for *V. vulnificus* (Tobit Regression, Wald-statistic: 5.917, p-value: 0.015), though for V. vulnificus only 3 datapoint had positive values for both oyster and water concentrations, thus further validation is needed (Additional File 3).

In addition to bacterial targets, we used qPCR to screen for the presence of *Ostreid herpesvirus 1* (OsHV-1) in whole oyster samples, which is a virus that commonly infects Pacific oysters causing mass mortality events. It has previously been detected in adult Pacific oysters in Southern California (29). We did not detect the virus in any of our samples.

### Influence of sample type on microbial community diversity and composition

We characterized microbial communities using 16S rRNA V4-5 amplicon sequencing for 4 different sample types: water, whole oysters, oyster gill tissue, and oyster digestive glands (DG). After filtering for quality, ~11M sequences were generated across samples and controls, representing ~33K unique amplicon sequence variants (ASVs) belonging predominantly to bacteria.

Sample type was a major driver of microbial diversity, with significant differences in both alpha and beta diversity. Water samples had higher species richness (Observed ASVs) than all oyster sample types, and water and whole oysters had higher alpha diversity (Shannon Diversity) than the oyster digestive gland and gill tissue (Figure 3A and 3B, Additional File 4). Sample type significantly influenced beta diversity (PERMANOVA, pseudo-F: 122.014, p-value: 0.001, Additional File 4), and sample types significantly differed from each other based on pairwise PERMANOVA tests (RPCA analysis, Figure 3C, Additional File 4).

**Figure 3:**
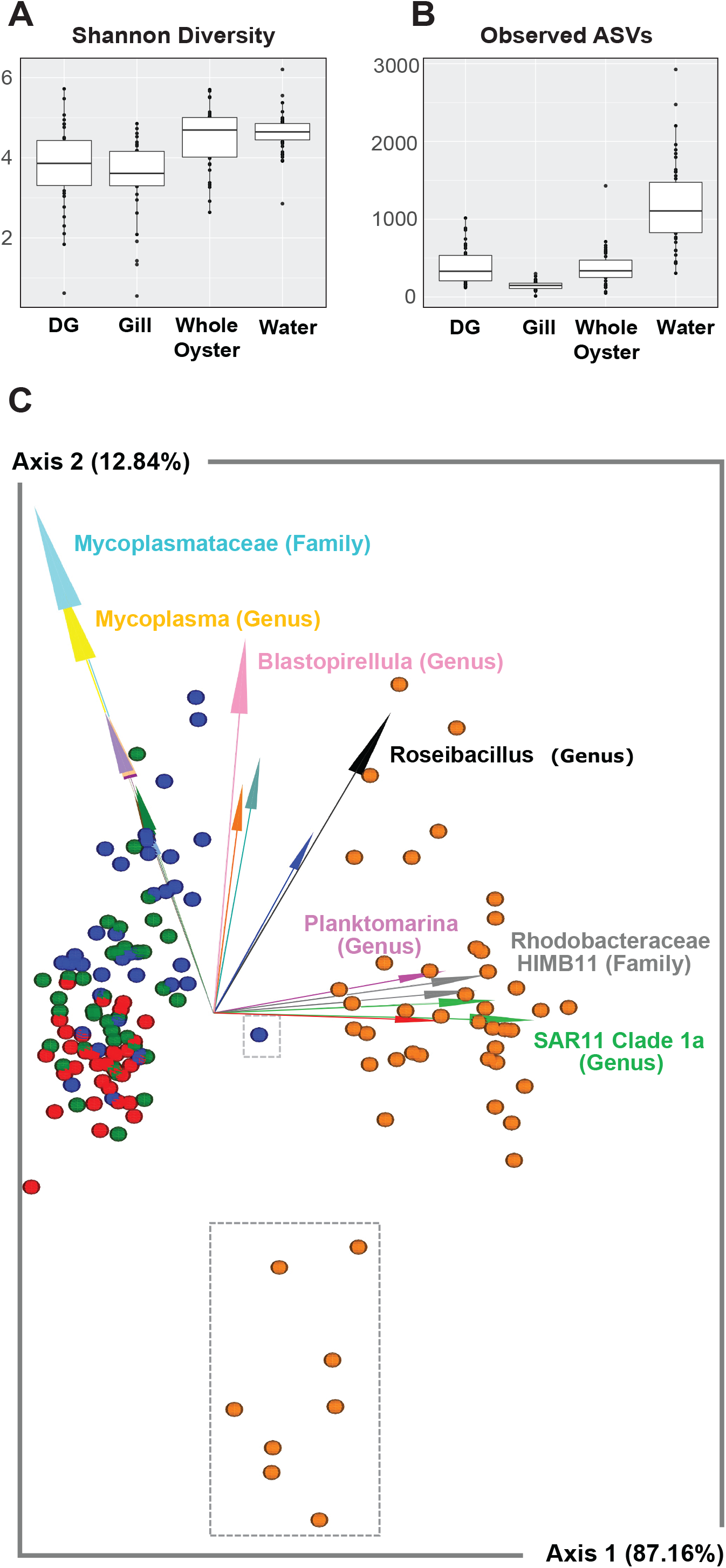
Diversity of water and oyster microbial communities and relationships with environmental parameters and pathogen concentrations. (A) Shannon diversity and (B) Observed ASVs compared between sample types. (C) DEICODE biplot showing RPCA distance between samples colored by sample type (blue = digestive gland, green = whole oyster, red = gill, and orange = water) and arrows depicting the key microbial taxa driving diversity, with annotation to the highest available taxonomic annotation noted. Water samples from sites NBS12 and NBS13 are enclosed in the larger gray dotted box, and a DG sample from NBS12 at time point 2 is enclosed in the smaller gray box.

Specific microbial taxa were strongly associated with either oyster or water sample types, or in some cases, specific oyster tissues. Key microbial features characteristic of oyster samples included members of the *Mycoplasmataceae* (Class Bacilli) and *Spirochaetaceae* families (Class Spirochaetia)(Figure 3C), which were also among the most relatively abundant taxa found in oyster tissues generally (Additional File 5, Additional File 6). Key water taxa included marine members of SAR11 *Clade Ia, HIMB11*, and *Planktomarina* (Figure 3C), which are all Alphaproteobacteria. While oyster gill and DG microbiomes shared some common microbial taxa, others were more representative of either sample type. For example, members of the genus *Blastopirellula* (Class Planctomycetes) and genus *Spiroplasma* (Class Bacilli) were more representative of DG samples, while *Sphingoaurantiacus* (Class Alphaproteobacteria), and Spirochaetia ASVs had high relative abundance in gill but not DG samples. (Additional File 6).

### Influence of site and environment on microbial community diversity and composition

Water microbial community diversity was impacted by site and environmental conditions to a greater extent than oyster microbiomes, which were comparably stable across sites. Beta diversity of water samples differed at back bay sites, which also exhibited differences in temperature, salinity, and chlorophyll *a* concentration (Figure 1). Specifically, microbial communities from NBS12 and NBS13 were significantly different from all other sites, and NBS11 was significantly different from all sites except 9 and 13 (Figure 3C, Additional Files 4 and 7). Communities from NBS12 and NBS13 were distinct from other sites on the biplot, as indicated by the grey dashed box surrounding these samples (Figure 3C), and in pairwise distance comparisons between sites within water samples (Additional File 7). Alpha diversity (Observed ASVs and Shannon Diversity) was negatively associated with temperature and positively associated with salinity in water samples, suggesting lower diversity at bay back sites (Additional File 8).

Beta diversity in oyster samples did not significantly differ across sites and environmental conditions, but there were site-specific differences in taxonomic composition and environmentally linked differences in tissue-specific alpha diversity. Whole oyster and DG samples had no statistically significant differences in alpha diversity across the variables tested; however, gill alpha diversity was associated with warmer temperatures (Additional File 8). Water and DG samples exhibited high relative abundance of one cyanobacterial ASV, which dominated microbial communities at Site 12 during weeks 2 and 4 (Additional File 5). The closest taxonomic assignment for this ASV was *Cyanobium* PCC-6307, which is commonly found in freshwater aquatic environments. Chlorophyll *a* levels were high during these sampling points which may suggest high actual and not just relative abundance of these cyanobacteria.

Additional differences in oyster microbial community taxonomy were observed at NBS12. Key taxa associated with DG microbiomes (Figure 4A, taxa annotated with arrows) had lower relative abundance at this site compared to the core DG microbiome (Figure 4B). Additionally, during week 2 in DG samples, a Chlamydiales ASV comprised ~30% of the microbial community. These are often intracellular animal pathogens and have been hypothesized to be linked to Oyster Oedema Disease (OOD) in pearl oysters (30). Both the *Cyanobium* and Chlamydiales ASVs had higher differential abundance in relation to the core DG microbiome at NBS12 compared to other sites (Figure 4C, 4D). The increase of these two ASVs at this time point also manifested as a community with distinct beta diversity (this data point is enclosed in a gray dotted box in the biplot in Figure 4C), however, this could not be statistically tested as oysters in this sample were pooled as a single data point.

**Figure 4:**
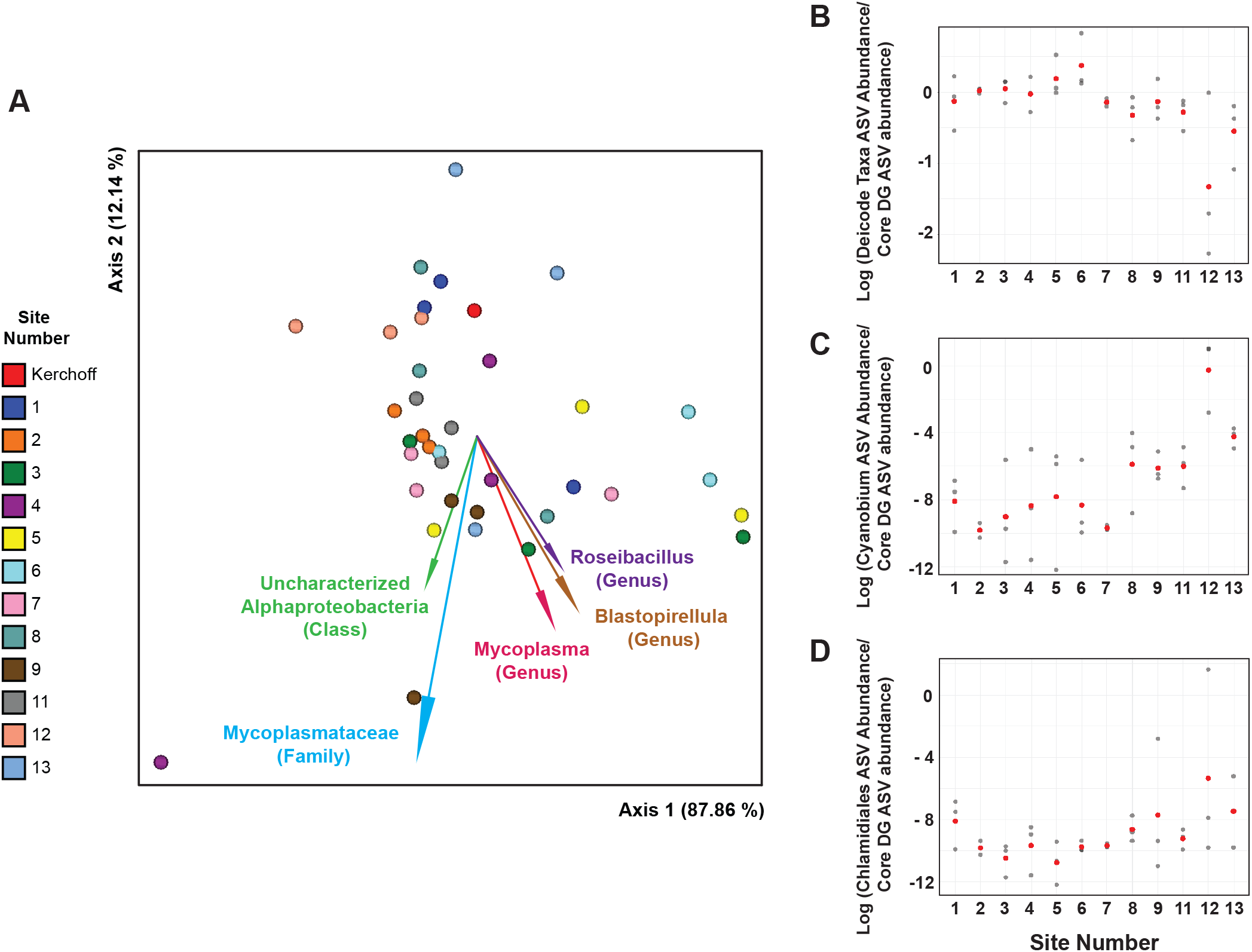
Differentially abundant taxa across sites in oyster digestive gland tissues. (A) RPCA biplot of digestive gland samples with markers depicting individual samples and marker colors indicating site (shown in legend). Arrows depict microbial taxonomic features driving differences in community composition (beta diversity) with the best available taxonomic composition. The log fold-change in differential abundance (compared as a ratio to the core digestive gland microbiome) is shown for (B) the driving taxonomic features from the RPCA DEICODE analysis (taxa with arrows), and two specific ASVs common at back bay sites: (C) Cyanobium PCC-6307, and (D) a Chlamydiales ASV. Red markers denote the average log-fold change.

### Relationship between potential bacterial pathogens and water or oyster microbial communities

We investigated whether certain characteristics of water and oyster microbial communities, namely community diversity and prominent microbial taxa in these sample types, were associated with high or low concentrations of fecal indicators and target *Vibrio* species.

In water samples, microbial communities with higher fecal indicator and *Vibrio* spp. levels were typically found at the back bay sites NBS12 and NBS13 and were significantly different in beta diversity compared to all other sites (Additional File 4, Additional File 7). Target bacteria concentrations coincided with lower salinity, warmer temperatures, and lower alpha diversity (Figure 1, Figure 3D). Both fecal indicator bacteria and potentially pathogenic *Vibrio* spp. were found in lower concentrations near the front bay sites and were significantly associated with higher salinities and linked to ASVs common to marine environments (e.g., SAR11 *Clade Ia, HIMB11*, and *Planktomarina*) (Figure 2, Figure 3, Additional File 9). Some specific microbial taxa were linked to low salinity conditions found at back bay sites and high potential pathogen concentrations (Additional File 9). For example, a *Flavobacterium* ASV was positively associated with multiple bacteria of concern, including elevated *Vibrio parahaemolyticus* (Vp) concentrations in both water and oysters. Additionally, *Cyanobium* PCC-6307 was abundant at sites and times when relatively high Vp levels were also observed (Site NBS12, weeks 2 and 4).

We also investigated whether the relatively high target bacteria concentrations observed in the water at sites NBS12 and NBS13, in environmental conditions that were potentially stressful for oysters, were also linked to changes in oyster microbial communities. As opposed to water samples, fecal indicator and *Vibrio* spp. concentrations in oyster samples didn’t exhibit clear trends except in the case of Vp, which was highest at back bay sites in both water and oyster samples (Figure 2). Similarly, beta diversity within oyster sample types did not vary significantly with site when grouped across weeks, though we did observe that the oyster DG microbial community where the highest concentration of Vp was observed (Week 2, Site NBS12) visibly separated from most DG samples in the DEICODE biplot (Figure 3C, gray dotted box).

Several oyster-associated microbial taxa were associated with either fecal indicator or *Vibrio* spp. concentrations in oysters. In whole oyster samples, *Mycoplasma* and *Blastopirellula* ASVs were negatively correlated with concentrations bacterial targets in water (Additional File 10). We also separately assessed gill and digestive gland (DG) microbiomes. In DG microbiomes, most of the high relative abundance taxa were not linked to environmental conditions or concentrations of human-health associated bacteria (Additional File 11). However, ASVs belonging to the *Blastopirellula* and *Mycoplasma* genera were negatively associated with *V. parahaemolyticus* in oysters, which likely drove the trend observed in whole oysters. Additionally, *Fuertia* bacteria (Order: Planktomycetales) were associated with lower levels of *E. coli* and Fecal Indicator bacteria. In Gill samples, most of the dominant taxa besides *Mycoplasma* ASVs were not significantly associated with environmental variables or human health-associated bacteria (Additional File 12). The cyanobacteria taxa *Cyanobium* PCC-6307 dominated water and oyster DG microbial communities where the highest concentrations of Vp were observed in oysters (Site NBS12, weeks 2 and 4), and where environmental conditions varied from other sites (Figure 1).

In oyster DG samples, *Cyanobium* PCC-6307 was positively associated with Vp concentrations in oysters, and positively associated with high concentrations of both the fecal indicators and *Vibrio* spp. quantified in water samples, meaning that when *Cyanobium* PCC-6307 was abundant in oyster DG microbiomes, potentially pathogenic bacteria concentrations were relatively high in the water and Vp was relatively high in the oysters (Additional File 11).

In addition to examining correlations between pathogen concentrations and relative abundance of microbial taxa in these sample types, we also used a differential abundance approach to assess whether other microbial taxa that were common in other DG or water samples decreased in relation to other taxa (i.e., a compositionally aware approach rather than solely examining relative abundance). This addressed the issue of whether the observed decrease in relative abundance of key taxa was due solely to the proportional increase in *Cyanobium* PCC-6307, or whether ratios of these taxa to the core DG microbiome increased as well. While the log ratio of *Cyanobium* to core DG taxa increased at site NBS12, the ratio of several key taxa of interest (as indicated by the DEICODE RPCA analysis and biplot, Figure 4A) decreased in relative abundance, suggesting that there was a decrease in the abundance of these taxa.

In investigating these associations between oyster microbiomes and fecal indicator and *Vibrio* spp. concentrations, some possible confounding variables in this dataset include the possibility that oyster microbiomes may change over time following translocation and that pathogens may take time to accumulate in oyster tissues. Specific properties of the sites where oysters were collected could influence oyster microbiomes after pooling, depuration, and deployment, and it is unknown how rapidly the oyster microbiomes stabilize and adapt to new conditions. Additionally, pooling multiple oyster samples, which is a common methodology for quantifying human pathogens, obscures variability of individual oysters in samples. Furthermore, the sites included in this study had varied physicochemical profiles. Although this allowed us to explore the effects of salinity and temperature, additional factors that were not measured may differ between sites and could influence the results as well. These present some limitations to the current experiment that should be considered.

## Discussion

### Oyster and water microbiota are differentially influenced by the environment

To investigate links between microbiomes and pathogen accumulation in bivalves, we first analyzed the microbial communities of water and examined their respective environmental influences. Though Pacific oysters originated in Japan, they are now extremely common along the North American West coast. However, the microbial communities and host interactions of these successful colonizers are poorly understood. Prior studies of regional Pacific oyster microbiomes observed similar microbial communities between extrapallial fluid, mantle fluid, and water microbial communities (34) and characterized impacts of different diets on fecal microbiomes (35). Our study focused on whole oyster as well as gill and digestive gland (DG) microbial communities, which are important to host physiology and may be prone to concentrating environmental bacteria since gills filter water particles and the DG may receive bacteria attached to food particles. All sample types we examined had significantly distinct microbial communities (Figure 3C, Additional File 5), which is consistent with prior studies showing that microbiomes of many animals, including oysters have tissue-specific microbial communities (36, 37).

Understanding how the environment influences oyster microbiomes is critical to preserving their ecosystem functions and supporting safe and abundant shellfish harvesting. We observed key taxonomic features among oyster sample types that may play a role in maintaining host health or reflect environmental adaptations. The dominant tissue-specific taxa we observed are common in Pacific oysters from diverse geographic regions, suggesting adaptation to oyster microenvironments and potentially host interactions. Furthermore, the tissue-specific communities were similar to each other across sites despite differing significantly from the surrounding water microbiomes. In particular, Spirochaetaceae and Mycoplasmataceae taxa were common in all three oyster sample types, with some ASVs more represented in specific sample types (Additional File 6).

Spirochaetaceae taxa were observed in *C. gigas* gill samples from the Wadden Sea, Tasmania and New South Wales (NSW), Australia (36–39). Members of this can cause human disease (e.g., Lyme disease and syphilis), but are also commonly associated with marine animals ranging from gastropods to corals, and provide beneficial host functions including nitrogen and carbon fixation (40–43). Mycoplasmataceae bacteria were common in DG samples, a pattern observed in above studies and in samples from France (44) and in Olympia oysters (*Ostrea lurida*) from the Northwest US (45). Interestingly, this was not a dominant DG microbial group in Pacific oyster DG samples from Mexico, however, this may be due to different analytical methods (46). Like Spirochaetaceae, several Mycoplasmataceae taxa are implicated in human and animal disease while other related taxa naturally reside in animal microbiomes with potentially symbiotic roles. A metagenomic study of Eastern oyster (*Crassostrea virginicus*) microbiomes identified a dominant *Mycoplasma* sp., and genomic and metabolomic reconstructions revealed a reduced set of metabolic functions and high reliance on host-derived nutrients (47). A reanalysis of previous *C. virginicus* from publicly available studies suggested that members of the parent Mollicutes class were highly prevalent in adult oyster DGs and lower in larva and biodeposits. For these taxonomic groups, their role in oysters or shellfish generally has not been characterized but may be of wide-spread importance, thus changes in the relative abundance of these taxa due to environmental or other stressors could have negative host impacts.

Community diversity of water strongly reflected the collection sites and associated environmental conditions, while in oyster samples beta diversity was more stable among sites and environments, but the relative abundance of key microbial taxa and alpha diversity differed. Variations in water microbial communities likely reflect diverse and dynamic coastal processes. Oysters, like many other animals, can to some extent regulate their microbial communities, though under stressful conditions these communities may shift from a stable to an unstable state (18, 48, 49). As our experimental sites represent a range of environmental conditions, particularly distinct at sites NBS12 and NBS13, we predicted that microbial communities would differ across sites. The lack of beta diversity divergence in oyster microbial communities deployed to sites with varying physicochemical water parameters and distinct water microbiomes (Figure 3C, Additional File 7) may reflect a strong selection pressure (host and/or microbe-mediated) for the structure of these communities, though it is also possible that short-term (i.e., weeks) exposure to these conditions does not impact community diversity, but that long-term exposure would. Alpha diversity was higher in gill (Observed ASVs and Shannon Diversity) and DG (Shannon Diversity) samples (Additional File 8) at the environmental conditions present at sites NBS12 and NBS13 (low salinity, high temperature), however, we did observe a negative association between these conditions and key microbial taxa present in DG tissues (Additional File 8), particularly members of the Mycoplasmataceae family discussed above, which could reflect a detrimental impact to hosts if these bacteria are beneficial.

### Potential human pathogen concentrations were linked to specific environments and microbial community features

Terrestrial and marine human pathogens often have distinct environmental abundance patterns based on their natural environments, but for both classes understanding associations with the marine environment and co-occurring microbes in water and oyster tissues may help characterize and ultimately predict bivalve pathogen concentrations. Some factors that likely play a role in bivalve pathogen accumulation include environmental pathogen concentrations, host behavior (e.g., feeding), pathogen interactions with bivalve transmission vectors, host physiology (including microbiome states), and microbial features that enable host colonization. Our study investigates these latter microbial variables, which are a key and understudied factor in human pathogen ecology in the marine environment.

Animal microbiomes are often linked to organismal health and the environment, and stressful environmental conditions can cause increases in the occurrence and relative abundances of otherwise uncommon bacterial taxa. In edible bivalve species, this may include an increase in human pathogens, but this possibility has not been adequately investigated. We predicted that oysters translocated to potentially stressful locations (i.e., sites NBS12 and NBS13) would concentrate more human pathogens, with coinciding microbial community changes. This occurred in the case of *Vibrio parahaemolyticus* (Vp), the most common cause of seafood transmitted vibriosis disease (4) but did not occur with regards to *V. vulnificus* (Vv) or fecal indicators measured, which exhibited variable patterns across environmental sites and over time. In water, FIB and *Vibrio* spp. bacteria were most abundant at sites NBS 12 and NBS13, consistent with their affinities for low salinity and warm temperatures. Fecal indicator and *Vibrio* spp. concentrations in oysters, however, were generally not linked to water concentrations or environmental conditions (except for Vp, discussed below), though in the case of Vv the low number of detections could confound the correlation analysis. It is possible that oysters with stable microbial communities may accumulate environmental human pathogens through feeding/filtering but possess microbiome-regulated controls on total accumulation. This would explain why, with the exception of Vp, oysters in water that contain relatively high Vv or FIB concentrations (i.e., the back bay sites NBS12 and NBS13) don’t accumulate proportionally more of these organisms than oysters at “low pathogen” sites. This also confirms that while environmental conditions conducive to pathogen proliferation may be necessary for high concentrations in water, they are not sufficient for accumulation in shellfish.

In the case of Vp, concentrations in water and oyster samples were positively correlated and linked to changes in oyster microbial communities, particularly at site NBS12, and to relatively warm temperature, low salinity environmental conditions. Prior studies have observed stress-induced increases and changes in *Vibrio* spp. relative abundance in oyster microbiota, but these studies typically focus on *Vibrio* spp. that are not human pathogens and typically do not quantify the bacteria (36, 50). The highest oyster Vp concentrations were observed at NBS12 during week 2, which featured a distinct DG microbial community (Figure 4C [small gray dotted box]). These high Vp concentrations coincided with an increase in the relative abundance of two uncommon DG taxa that could suggest an unhealthy host state: a Chlamydiae family ASV, which are often intracellular animal pathogens and potentially cause Oyster Oedema Disease (OOD) in pearl oysters (30), and a cyanobacteria taxon *Cyanobium* PCC-6307, which was commonly present in both water and oyster DG samples (Additional File 5, Figure 4B, 4C). Meanwhile, several common DG oyster taxa (e.g., *Mycoplasma* and *Blastopirellula*) were less relatively abundant at the “high Vp” back bay sites. Since *Mycoplasma* bacteria are considered core members of oyster DG microbial communities and have been linked to increased oyster survival (45) the increase in rare taxa and decrease in core taxa may suggest poor host health and microbial community dysbiosis. Despite these observations at NBS12, Vp was also consistently abundant at site 13, which had similar environmental conditions but not these divergent taxa. This may suggest that both the environment and specific microbial taxa in oysters and the environment act synergistically to enable marine human pathogen accumulation in oysters. Other factors may also be involved, including time oysters are exposed to stressful environments and oyster feeding behavior.

### Phytoplankton links to oyster pathogen accumulation: signs of dysbiosis or transport vectors?

The highest concentration of *Vibrio parahaemolyticus* bacteria we observed in oysters, which causes an estimated 45,000 cases of vibriosis annually in the US (4), was positively linked to elevated relative abundance of a cyanobacteria ASV in both water and oyster DG samples. This led us to consider whether the abundance of this taxon signaled dysbiosis in oyster DG microbiomes, whether cyanobacteria were acting as a transport vector for oyster pathogen accumulation (via feeding), or potentially both. Chlorophyll *a* concentration was positively associated with Vp (Figure 3D, 3E) and was also highest at NBS12 compared to samples collected the same week at other sites, which coincided with the observed cyanobacterial taxa (Additional File 6). This supports the possibility that these cyanobacteria were abundant in the microbial community, which would increase their likelihood of interacting with Vp, though we did not measure actual attachment and other phytoplankton taxa may produce chlorophyll *a* in the environment.

Since oysters are filter-feeders that consume particles and phytoplankton prey, positive associations between pathogens and aquatic taxa such as *Cyanobium* spp., could indicate vibrio attachment mediating oyster pathogen accumulation. Vp attachment to phytoplankton could enable environmental persistence by providing nutrients and attachment substrate while increasing oyster accumulation if they are subsequently preyed upon, as both terrestrial and marine pathogens frequently attach to marine particles and form highly concentrated biofilms. We previously identified phytoplankton taxa positively linked to high Vp concentrations in southern California water microbiomes, including common oyster prey species (e.g., *Thalassiosira* spp. diatoms) and a cyanobacterial ASV *Prochlorothrix* sp. PCC-9006 (13), though we did not assess oyster pathogen concentrations in that study.

It is unclear whether Vp actually attaches to cyanobacteria in the environment, and whether cyanobacteria could serve as a vector for Vp uptake by oysters. Vp interactions have been demonstrated in a laboratory setting for cyanobacteria and other phytoplankton taxa (15, 51), and attachment would be consistent with the widespread particle-attaching lifestyle of *Vibrio* species. While cyanobacteria are often small and not known to be a major bivalve food source, *Cyanobium* spp. can form larger cell aggregates (52), which would enhance their potential for both vibrio attachment and oyster uptake. Given the other changes observed in DG microbiomes that coincided with high Vp abundance, cyanobacteria abundance may also represent transient environmental bacteria accumulating in oysters due to altered host microbiome states and potentially impaired host health. Our observation that alpha diversity was higher in DG samples at these Back Bay sites may suggest hosts were not actually impaired since lower alpha diversity is often linked to impaired host states. However, recent studies have called this paradigm into question with observations that alpha diversity is not necessarily linked to host health (53, 54). A recent study of Eastern oysters observed no significant differences in alpha diversity between infected and uninfected individuals for most tissue types, including gut microbial communities (47). Furthermore, microbiome diversity is far from the sole indicator of host stress. While we are not able to discern from our study which of these scenarios occurred, or if it was a combination of factors, this represents an important hypothesis to test in future studies.

Several future research directions can further strengthen our understanding of human pathogen ecology in the marine environment and enable safe seafood consumption now and in future ocean conditions. Investigating changes in the eukaryotic microbiota may reveal additional vibrio transmission vectors and microbiome dynamics linked to host health not clear from bacterial microbiota analyses. Integrating knowledge of the oyster microbiome with host physiology, particularly feeding behavior, will help elucidate the interactions between both host and microbe-mediated drivers of bivalve pathogen accumulation, particularly in relation to predicted future ocean changes. This could also include assays to observe the actual attachment of pathogenic *Vibrio* spp. to environmental microbes and oyster tissues (e.g., FISH (55). Furthermore, better understanding the microbial taxa either positively or negatively linked to oyster pathogen accumulation is essential to characterizing the ecology of these infectious disease organisms and could ultimately lead to developing bioindicators of oyster pathogen accumulation or identifying taxa that could be used as defensive probiotics.

## Conclusion

Understanding factors beyond environmental conditions that drive human pathogen accumulation in bivalves is critical to supporting safe and sustainable seafood consumption. Here we demonstrate that both environmental factors and microbial communities interact to differentially influence concentrations of potential human pathogens in water and oysters. While specific environmental conditions are linked to both microbial community diversity and concentrations of potential pathogenic *Vibrio* spp. and indicator species indicative of fecal contamination in water samples, concentrations of these bacteria in water are not necessarily linked to concentrations in oysters. Oyster microbiomes and pathogen concentrations were less environmentally dependent than those in water, except in the case of *Vibrio parahaemolyticus*, where relatively high oyster concentrations were associated with an increase in environmental cyanobacteria and a decrease in the relative abundance of key digestive gland taxa. This study suggests that environmental conditions and microbial communities may interact, potentially synergistically, to drive human pathogen concentrations in oysters. Future research integrating pathogen attachment to oyster uptake vectors in the environment, oyster behavior and physiology, and the functional roles of oyster microbiomes and specific taxa - particularly in response to changing environmental conditions - will provide data critical for promoting safe seafood harvesting for a growing human population in the future.

## Supporting information

Additional Files for Diner et al. 2022

## Acknowledgements

We acknowledge Rachel Noble and Brett Froelich for project guidance and assistance in culturing, identifying, and screening *Vibrio* species bacteria. Funding was provided to R.E.D. by a San Diego Institutional Research and Academic Career Development Award (IRACDA) (NIH/NIGMS IRACDA K12 GM068524) and the National Science Foundation postdoctoral research fellowship in biology (PRFB; P2011025). This study was supported, in part, by National Science Foundation (NSF-OCE-1637632 and NSF-OCE-1756884), National Oceanic and Atmospheric Administration (NOAA) (NA15OAR4320071 and NA19NOS4780181), and Gordon and Betty Moore Foundation (GBMF3828) grants to A.E.A.

## Author Contributions

R.E.D., A.Z-F., E.C., J.G., and A.E.A. were responsible for the conception and design of the study. R.E.D., A.Z-F., and E.C. were responsible for collecting and processing samples. R.E.D., A.Z-F., S.A., S.M.K., E.K., and Y.G. conducted data analysis. R.E.D., A.Z-F., S.A., and J.G. contributed to data interpretation and manuscript preparation.

