## Additional Files for Diner et al. 2022 for "Host and Water Microbiota are Differentially Linked to Potential Human Pathogen Accumulation in Oysters"

**Additional File 1: Molecular quantification methods for fecal indicator bacteria and target *Vibrio* spp.**

| Target Bacteria | Matrix | Method | Method Reference | Gene Target | PCR Reference |
|---|---|---|---|---|---|
| Fecal coliform/ *Escherichia coli* | Oyster | 5-tube, 3 dilution MPN MTF (LST, EC-mug for confirmation) | APHA, 1970 | | |
| Fecal coliform/ *E. coli* | Water | EPA Method 1603: Membrane filtration with mTEC | USEPA, 2009 | | |
| *Enterococcus* spp. | Water | EPA Method 1600: Membrane filtration on mEI | USEPA, 2009 | | |
| *Vibrio parahaemolyticus* | Oyster/Water | Spread plate on CHROMagar Vibrio + isolate confirmation by PCR | Froelich et al., 2017 | *toxR* | Taiwo et al. 2017 |
| *Vibrio vulnificus* | Oyster/Water | Spread plate on CHROMagar Vibrio + isolate confirmation by PCR | Froelich et al., 2017 | *vvhA* | Warner and Oliver, 2008 |

**Additional File 2:** Metadata file containing data used for community analysis and raw data pertaining to experimental sites, environmental conditions, and pathogen concentrations.

Available for download on Figshare: https://doi.org/10.6084/m9.figshare.21272916

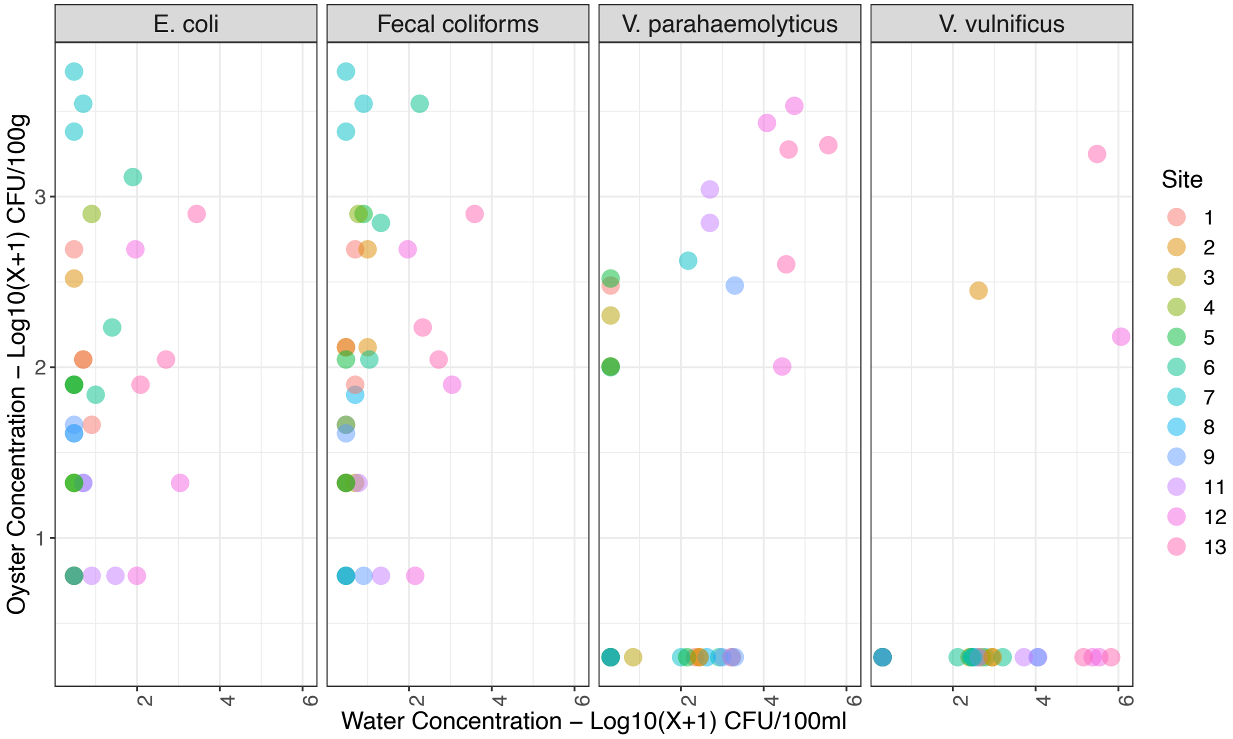


**Additional File 3: Relationships between water and oyster concentrations of fecal indicator bacteria (FIB) and target *Vibrio* species.** FIBs include *Escherichia coli* and fecal coliform bacteria. *Vibrio* spp. targets include *V. parahaemolyticus* and *V. vulnificus*. Correlations between water and oyster concentrations were not significant for *E. coli* and fecal coliforms (Linear Regression, *E. coli*: F-statistic: 0.6927, adjusted p-value: 0.411, fecal coliform: F-statistic: 1, adjusted p-value: 0.1916). Correlations between water and oyster concentrations were significant for *V. parahaemolyticus* (Tobit Regression, Wald-statistic: 5.862, p-value: 0.015) and for *V. vulnificus* (Tobit Regression, Wald-statistic: 5.917, p-value: 0.015), though for *V. vulnificus* only 3 datapoint had positive values for both oyster and water concentrations.

**Additional File 4: Alpha and Beta Diversity Statistics**

Available for download on Figshare: https://doi.org/10.6084/m9.figshare.21273009

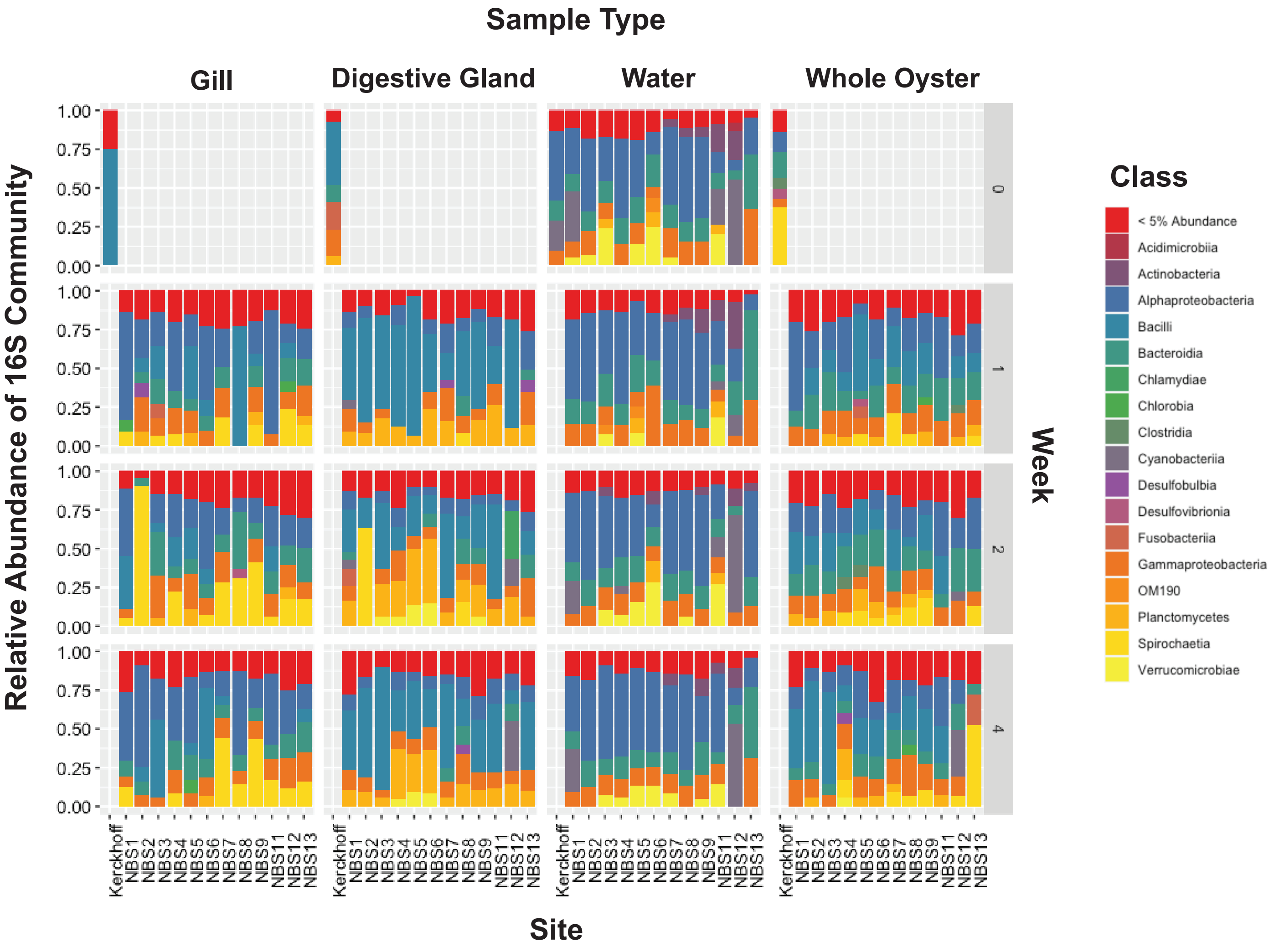


**Additional File 5: Taxonomic composition of oyster and water microbial communities over time.** Relative abundance of 16S microbial communities for all sample types (water, whole oyster, gill, and digestive gland) at each week and site, including the Kerchoff facility following depuration (pre-deployment). Taxa are presented by class and classes representing less than 5% of the sequences in a particular sample type are grouped together (depicted by the red <5% category).

**Additional File 6: Top 10 most relatively abundant ASVs in water and oyster tissue samples with genus-level taxonomic annotation.**

Available for download on Figshare: https://doi.org/10.6084/m9.figshare.21273054

**Additional File 7: Pairwise differences in beta diversity between sites for each sample type, based on RPCA analysis.** Box plots and statistics can be viewed and downloaded by downloading and viewing the following visualization files for each sample type on https://view.qiime2.org/ (29):

**Water:** NBS19_16S_autoRPCA_Water_Site_significance.qzv

**Whole Oyster:** NBS19_16S_autoRPCA_WholeOyster_Sitenumber_significance.qzv

**Digestive Gland:** NBS19_16S_autoRPCA_Gut_Sitenumber_significance.qzv

**Gill:** NBS19_16S_autoRPCA_Gill_Sitenumber_significance.qzv

Available for download on Figshare: https://doi.org/10.6084/m9.figshare.21273099

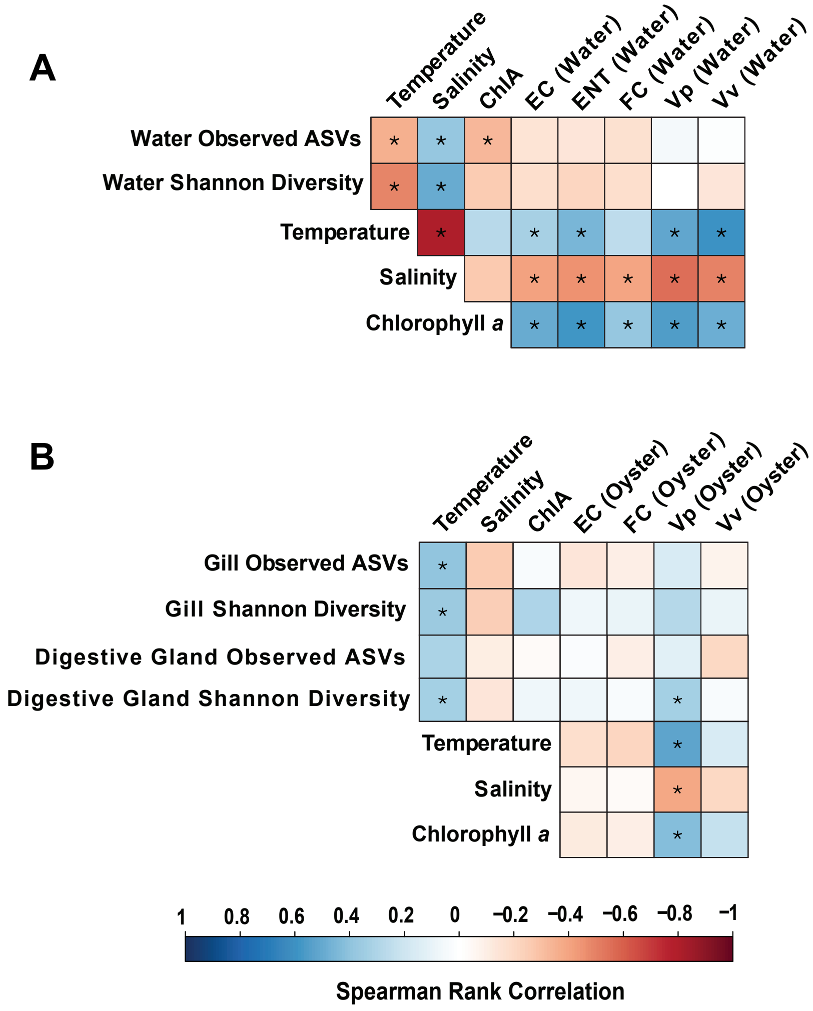


**Additional File 8: Spearman rank correlations of alpha diversity metrics, environmental variables, and target bacteria concentrations.** Observed ASVs and Shannon Diversity alpha diversity metric correlations are shown for (A) water samples and (B) oyster tissue samples. Blue squares indicate positive correlations while red squares indicate negative correlations between variables.

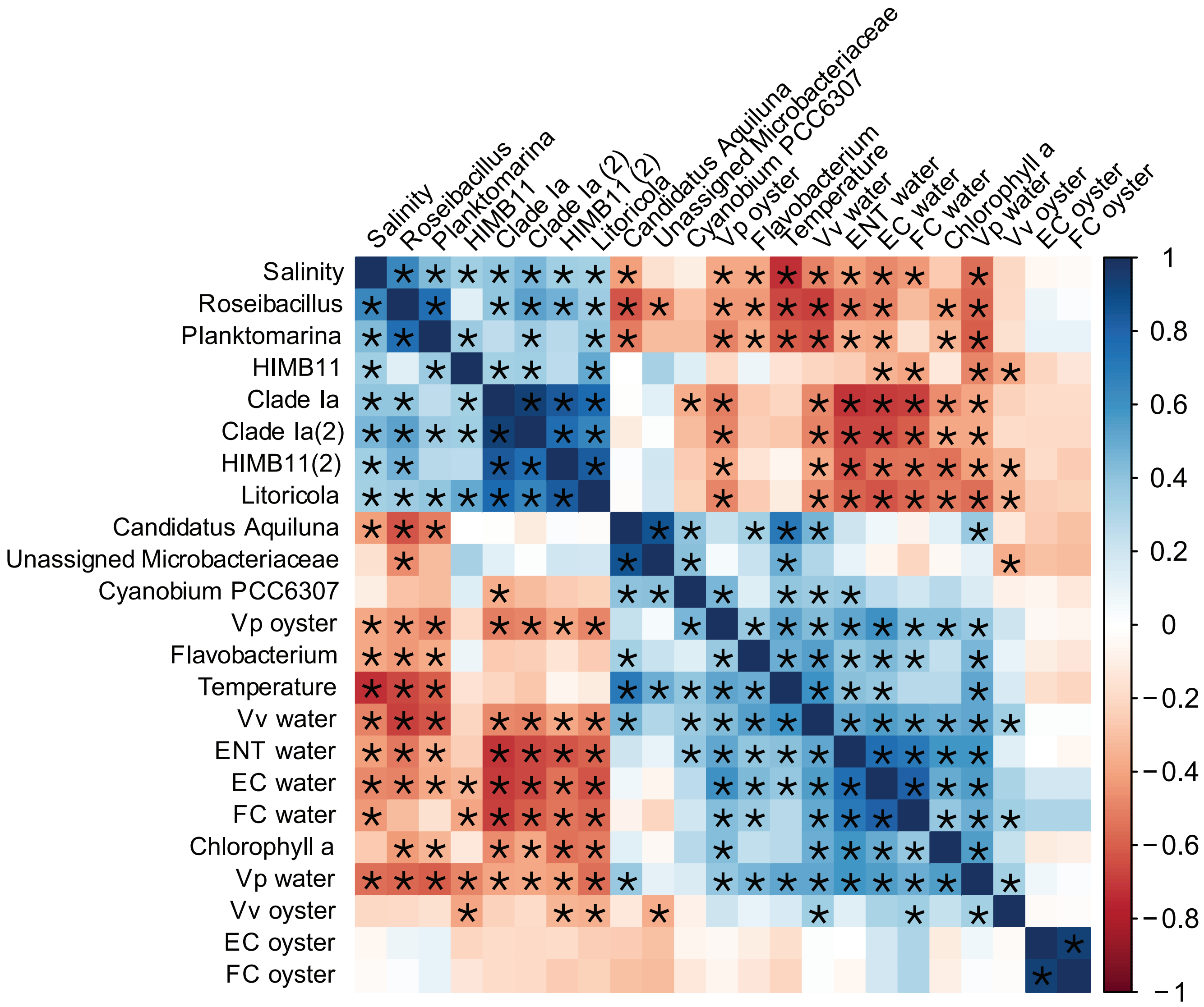


**Additional File 9: Spearman Rank correlations between fecal and vibrio bacteria concentrations, environmental variables, and the top 10 most abundant ASVs and specific bacteria of interest in <u>water</u> samples.** Environmental variables (temperature, salinity, and chlorophyll *a*), and target bacteria concentrations are quantitative values, while microbiome taxa of interest are relative abundance. Bacterial taxa are annotated to the best available taxonomic resolution and are further described in Additional File 6. Numbers in parentheses indicate multiple ASVs with the same annotation. Target bacteria abbreviations for water and oysters are: EC = *Escherichia coli*, FC = fecal coliform bacteria, Vp = *Vibrio parahaemolyticus* and Vv = Vibrio vulnificus. * denotes a significant correlation (p-value = <0.05), and blue squares indicate a positive correlation while red squares indicate a negative correlation.

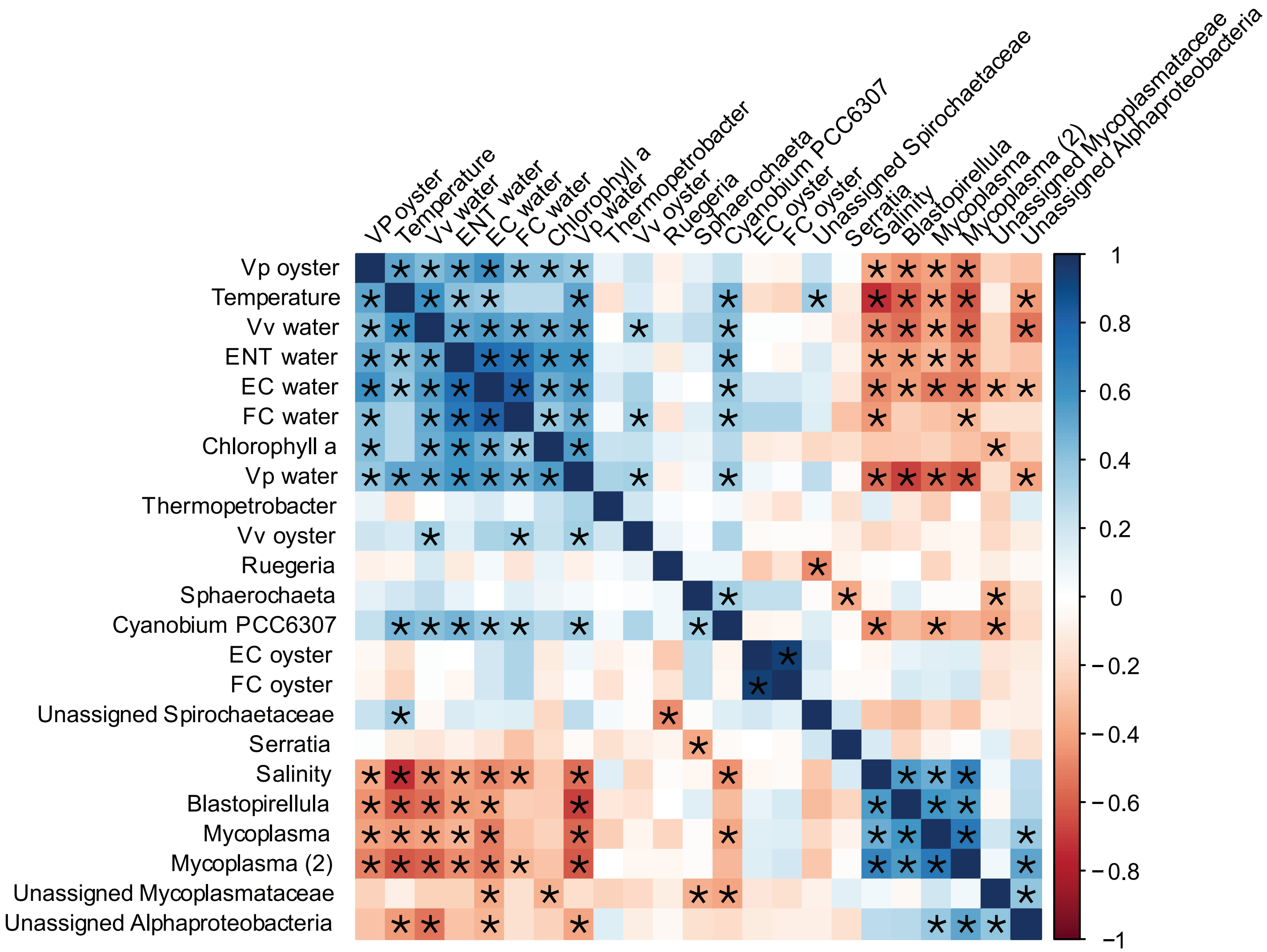


**Additional File 10: Spearman Rank correlations between fecal and vibrio bacteria concentrations, environmental variables, and the top 10 most abundant ASVs and specific bacteria of interest in <u>whole</u> oyster samples.** Environmental variables (temperature, salinity, and chlorophyll *a*), and target bacteria concentrations are quantitative values, while microbiome taxa of interest are relative abundance. Bacterial taxa are annotated to the best available taxonomic resolution and are further described in Additional File 6. Numbers in parentheses indicate multiple ASVs with the same annotation. Target bacteria abbreviations for water and oysters are: EC = *Escherichia coli*, FC = fecal coliform bacteria, Vp = *Vibrio parahaemolyticus* and Vv = Vibrio vulnificus. * denotes a significant correlation (p-value = <0.05), and blue squares indicate a positive correlation while red squares indicate a negative correlation.

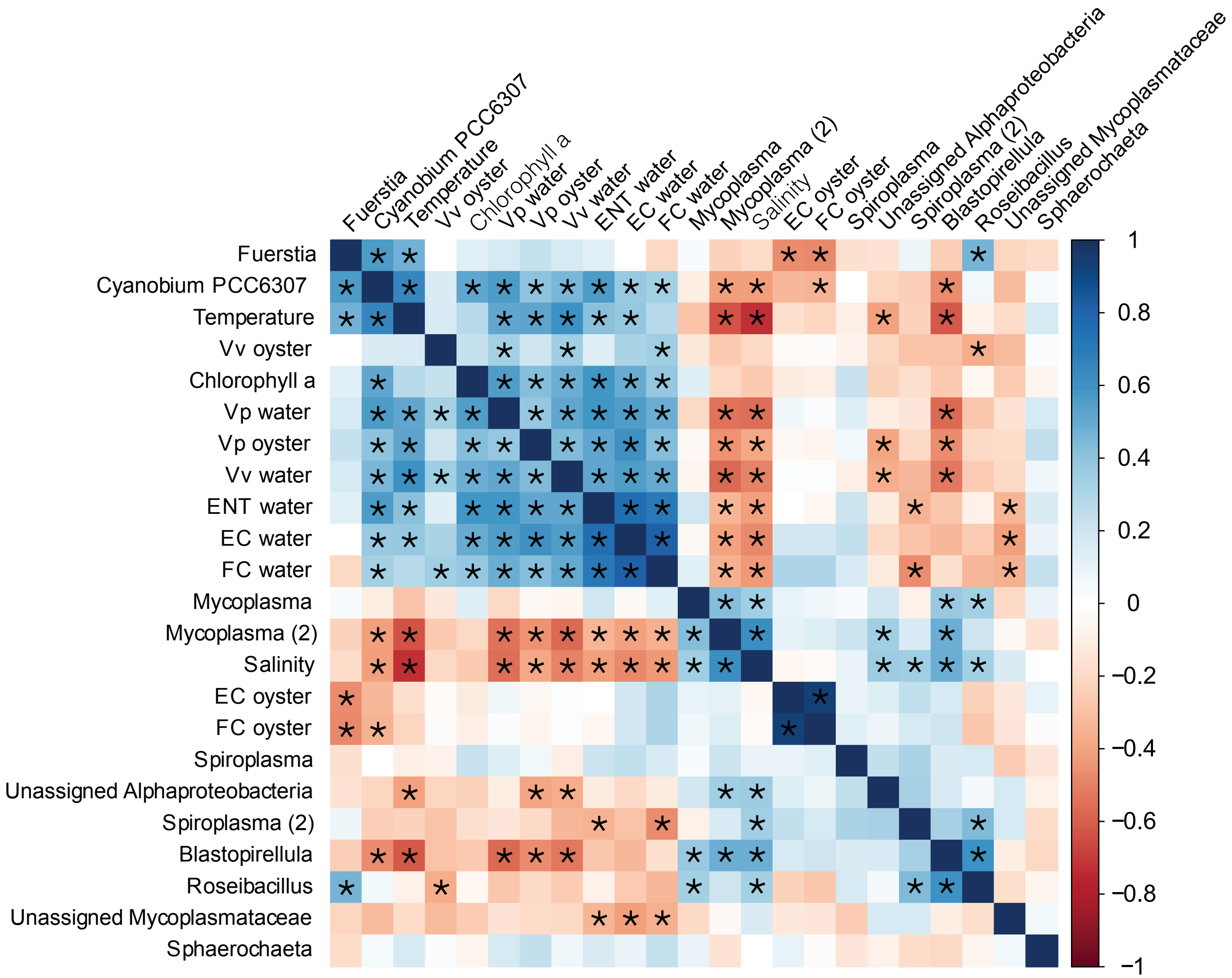


**Additional File 11: Spearman Rank correlations between fecal and vibrio bacteria concentrations, environmental variables, and the top 10 most abundant ASVs and specific bacteria of interest in digestive gland samples.** Environmental variables (temperature, salinity, and chlorophyll *a*), and target bacteria concentrations are quantitative values, while microbiome taxa of interest are relative abundance. Bacterial taxa are annotated to the best available taxonomic resolution and are further described in Additional File 6. Numbers in parentheses indicate multiple ASVs with the same annotation. Target bacteria abbreviations for water and oysters are: EC = *Escherichia coli*, FC = fecal coliform bacteria, Vp = *Vibrio parahaemolyticus* and Vv = Vibrio vulnificus. * denotes a significant correlation (p-value = <0.05), and blue squares indicate a positive correlation while red squares indicate a negative correlation.

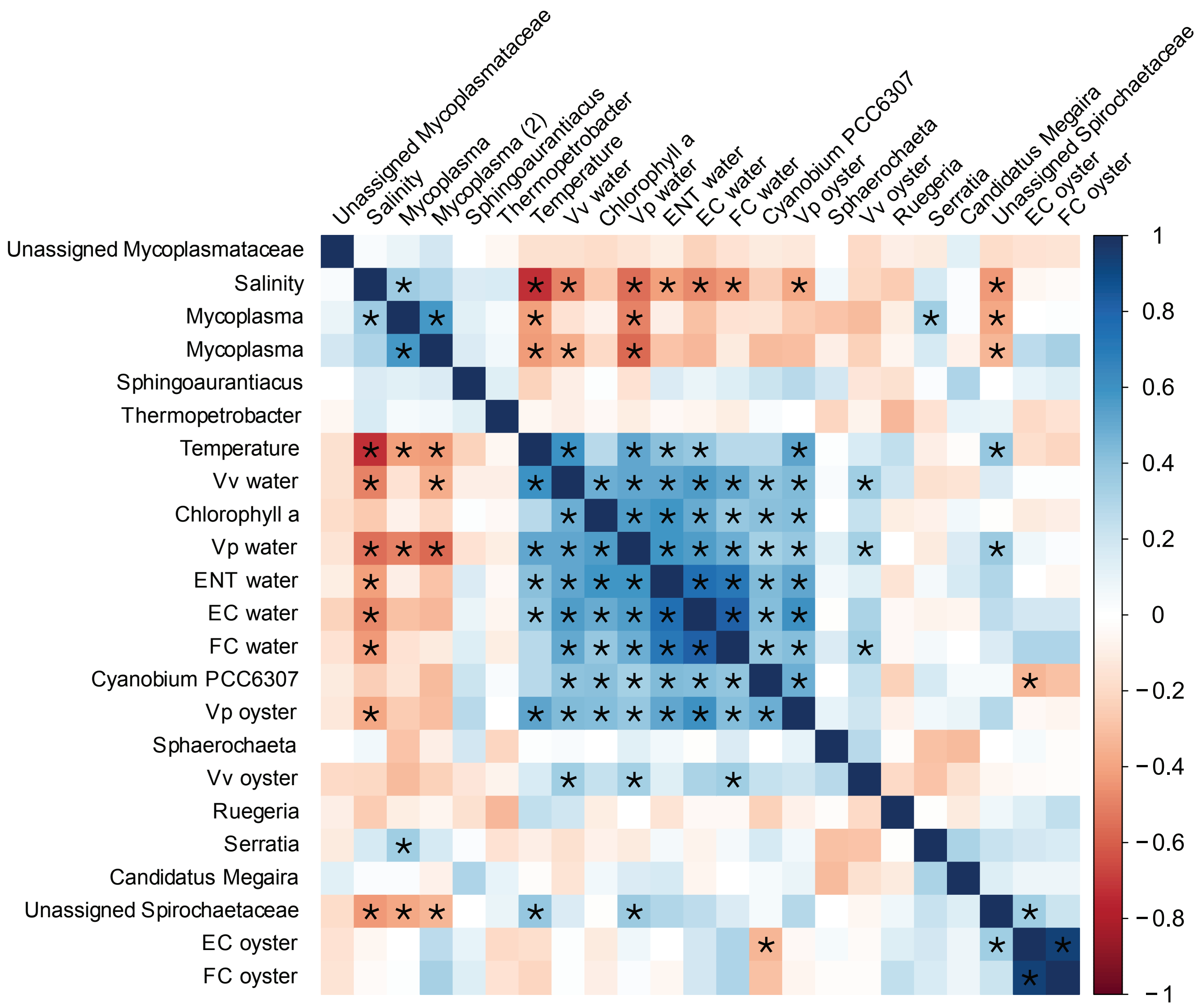


**Additional File 12: Spearman Rank correlations between fecal and vibrio bacteria concentrations, environmental variables, and the top 10 most abundant ASVs and specific bacteria of interest in gill samples.** Environmental variables (temperature, salinity, and chlorophyll *a*), and target bacteria concentrations are quantitative values, while microbiome taxa of interest are relative abundance. Bacterial taxa are annotated to the best available taxonomic resolution and are further described in Additional File 6. Numbers in parentheses indicate multiple ASVs with the same annotation. Target bacteria abbreviations for water and oysters are: EC = *Escherichia coli*, FC = fecal coliform bacteria, Vp = *Vibrio parahaemolyticus* and Vv = Vibrio vulnificus. * denotes a significant correlation (p-value = <0.05), and blue squares indicate a positive correlation while red squares indicate a negative correlation.
